# Theta-rhythmic attentional exploration of space

**DOI:** 10.1101/2025.08.16.670674

**Authors:** M. Senoussi, L. Galas, N.A. Busch, L. Dugué

**Author notes:** shared first authorship (M. Senoussi, L. Galas). Corresponding author: Mehdi Senoussi –.

## Abstract

Attention facilitates stimulus processing by selecting specific locations (spatial attention) or features (feature-based attention). It can be sustained on a given location or feature, or re-oriented between locations or between features, enabling attentional exploration. Sustained attention was associated with alpha (8-12Hz^1–4^) oscillations. More recently, authors suggested that exploratory attention was instead related to theta (4-7Hz; for review^5–7^). To date, there is no systematic evaluation of attentional exploration across stimulus dimensions (space, features) in relation to theta oscillations. Using attentional cueing and electroencephalography (EEG) in humans, we assessed exploration of stimulus dimension during the (1) precue-to-stimulus (first attentional orienting) and (2) post-stimulus (during stimulus processing) trial periods. In the precue-to-stimulus period, we classified the cued dimension (attending to a location/feature) from EEG topographies to assess the neural dynamics of sustained and exploratory attentional orienting. Temporal generalization matrices showed oscillatory patterns of classification accuracies across time. When attention was sustained on a cued location but explored the feature dimension as the target feature was unknown, only the alpha frequency was observed. When attention was sustained on a cued feature but explored locations, both alpha and theta frequencies were observed. Focusing on post-stimulus theta oscillations revealed increased theta power in invalid trials (attention reorients from the precued distractor location/feature to the target –exploration) relative to valid trials in both feature-based and spatial attention conditions. Post-stimulus theta oscillations further predicted behavioral performance. Together, our results show that while alpha oscillations are associated with sustained attention regardless of the attended dimension, theta oscillations are specifically related to spatial exploration.

## Results

Humans constantly receive an overwhelming amount of sensory information. Attention is the cognitive function able to allocate resources to the efficient processing of task relevant stimuli^8^. Attention can be sustained at a spatial location or on a stimulus feature and influence perceptual performance for up to several seconds^9^. Spatial and feature-based attention can also explore by switching from a stimulus or feature to another (for review^5–7,10^).

Alpha oscillations have been systematically reported when both spatial and feature-based attention are sustained. Specifically, the topography of alpha oscillations lateralizes^1–4^, with stronger alpha power at the ipsilateral electrodes relative to the attended location in the visual field. In the case of spatial attention, this lateralization occurs in the precue-to-stimulus period, i.e., during the sustained attentional orientation toward a specific location. For feature-based attention, the location of the attended feature is not known in the precue-to-stimulus period, however, in the post-stimulus period it is, and lateralization was observed^11^.

The link between exploratory attention and low frequency oscillations is, however, contested. Over the last two decades researchers have proposed that attention can explore the visual scene periodically at the theta frequency, and that this sampling is supported by brain oscillations at corresponding frequencies^5–7,10^. In visual search tasks, for instance, in which participants look for a target stimulus among distractors, the pre– and post-stimulus phase of theta oscillations predict successful target discrimination^12–15^, and behavioral performance is periodically modulated at the same frequency^12,16–19^, for review^5^. This attentional sampling of the visual space was observed in several other studies using a cueing manipulation of several attention types, including voluntary and involuntary spatial, feature– and object-based attention (e.g.,^20–24^), and was sometimes related to corresponding neural rhythms (e.g.,^25,26^). These results suggest a link between theta oscillations and exploratory attention regardless of the attended dimension. However, this conclusion has not been systematically studied as it was drawn from a set of independent studies. Here, we explicitly test this proposal by manipulating the attended stimulus space (location/feature) within the same study and participants.

### Spatial and Feature-based attentional cueing

Voluntary (endogenous) Spatial Attention (SA) and Feature-Based Attention (FBA) were manipulated in two separate experimental sessions in which EEG was concurrently recorded (**Figure 1A**). Behavioral performance (**Figure 1B**) was evaluated using two 2×2-factor repeated measurement ANOVAs (validity x attentional dimension, separately for d-primes and reaction times). Reaction times were faster and d-primes were higher for valid compared to invalid trials (d-prime: F(1,15)=17.21, p<0.001, generalized eta squared, g-η²=0.099; reaction times: F(1,15)=9.46, p=0.008, g-η²=0.126), but there was no effect of attention dimension (d-prime: F(1,15)=0.54, p=0.473, g-η²=0.009; RT: F(1,15)=0.95, p=0.344, g-η²=0.003) and no interaction (d-prime: F(1,15)=0.83, p=0.375, g-η²=0.005; RT: F(1,15)=0.07, p=0.790, g-η²<0.001). Together, the results show that both types of attention were successfully manipulated (and with similar overall performance), with no speed-accuracy tradeoffs^23,27–32^.

**Figure 1.**
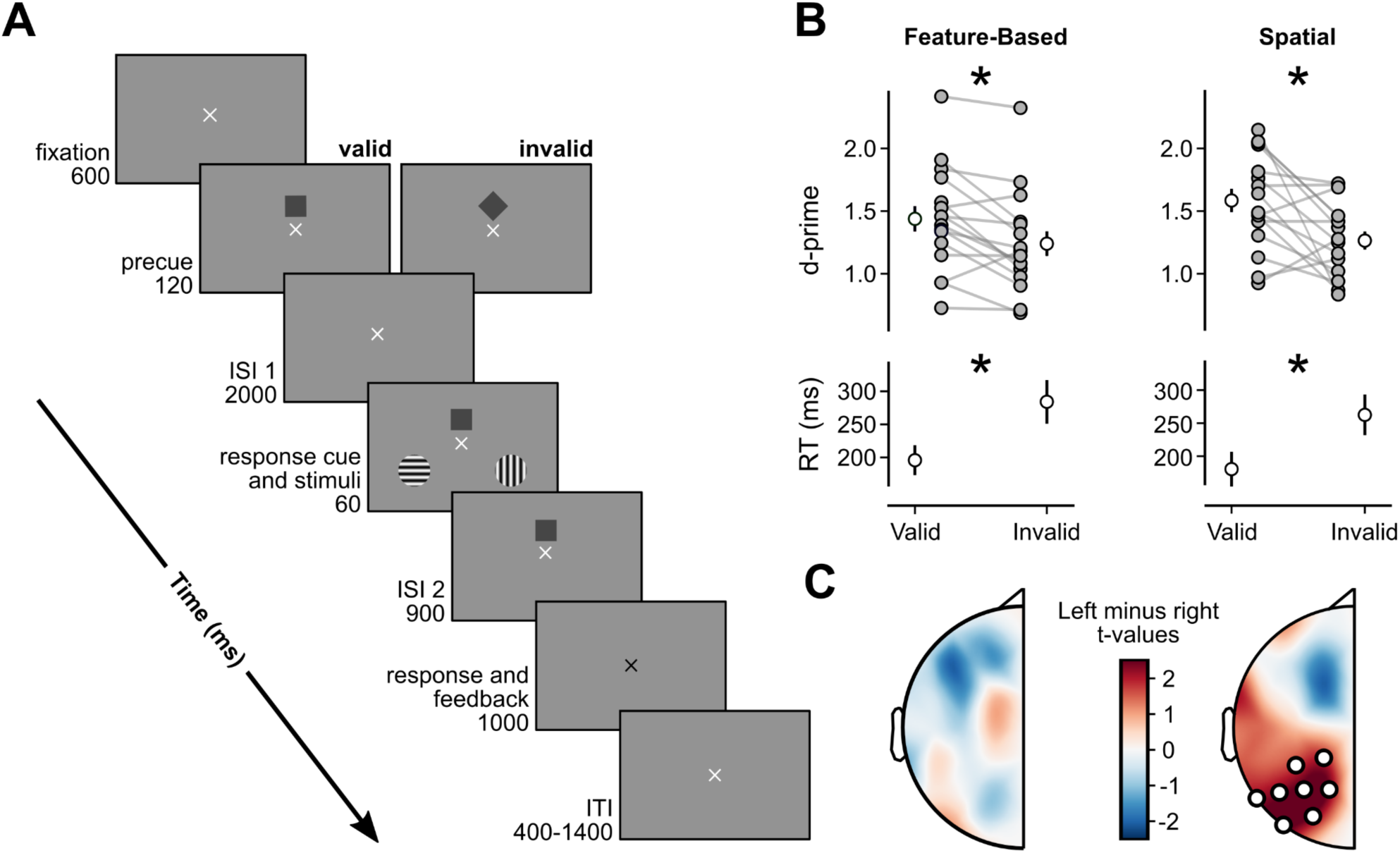
Experimental design and manipulation assessment. **A.** Trial sequence. Trials started after 600 ms of fixation. Then the precue was displayed for 120 ms instructing the participant to attend to a specific location or feature (they learned the association between each cue, square/diamond and the stimulus dimension in a previous training session). After a first ISI of 2000 ms, a response cue and two gratings were displayed for 60 ms. The response cue indicated which one of the two gratings was the target and was 75% of the time the same as the precue (valid trials). The response cue stayed 900 ms (ISI 2) longer on the screen and then participants had to report the tilt of the target (clockwise or counterclockwise around the vertical or horizontal axis). **B.** Behavioral results. Individual and averaged d-prime for spatial and feature-based attention in valid and invalid conditions. * indicate significant differences (two-way repeated measure ANOVAs, d-prime: p<0.001, RT: p<0.008). **C.** Alpha lateralization maps. T-values of the Left minus Right difference in alpha spectral power estimates for signal extracted from 500 to 1000 ms from cue onset (positive values indicate an increase in alpha amplitude in the ipsilateral region to the attended location). White dots: electrode within a cluster of significant positive difference to zero (permutation cluster test, SA: p = 0.022)

We further tested whether alpha lateralization, i.e., increase in alpha power in ipsilateral electrodes relative to the cued location, was present in SA but not in FBA, thus replicating previous research^4,11^. In SA, we subtracted alpha power in attend-left trials from attend-right trials. In FBA, we subtracted alpha power in attend-to-vertical from attend-to-horizontal trials. While in the SA condition we observed a significant cluster of electrodes exhibiting the expected alpha lateralization (p=0.022; **Figure 1C**), there was no lateralization in FBA (p > 0.05; comparing FBA trials in which targets appeared on the left versus right led to the same results). These results together with the behavioral performance confirm expected results from a successful attentional manipulation.

### Decoding attentional orienting across stimulus space

To assess the dynamics of attentional orienting in stimulus space, we tested whether the attended location (SA) or feature (FBA) could be decoded from brain activity topographies. We performed a temporal generalization classification analysis on the signal from all electrodes in the precue-to-stimulus period (from –200 ms to 2,100 ms relative to precue onset; **Figure 1A**). The objective here is to analyze how classification of brain activity topographies is changing over time. We trained a classifier at a time point (t_i_) and tested it at every time point (t_j_) in the interval. The classification accuracy reflects whether the multivariate pattern of brain activity at time point t_i_ permits to decode the attended location or feature at this time point (when t_i_=t_j_), and whether this ability to discriminate generalizes to other time points (when t_i_≠t_j_; further details in **STAR Methods**). This approach allowed us to go beyond correlation between oscillations and task conditions, and evaluate the information content (i.e., the attended stimulus dimension) of neural activity. For both FBA and SA, we observed a significant cluster of above chance classification accuracy (FBA: p=0.0043, **Figure 2A**; SA: p=0.0001, **Figure 2C**). Both clusters encompassed a large portion of the analysis window, starting approximately at 100 ms from precue onset, and extending to the end of the precue-to-stimulus period. Due to the properties of cluster tests one cannot interpret the overall shape of the clusters any further^33^.

**Figure 2.**
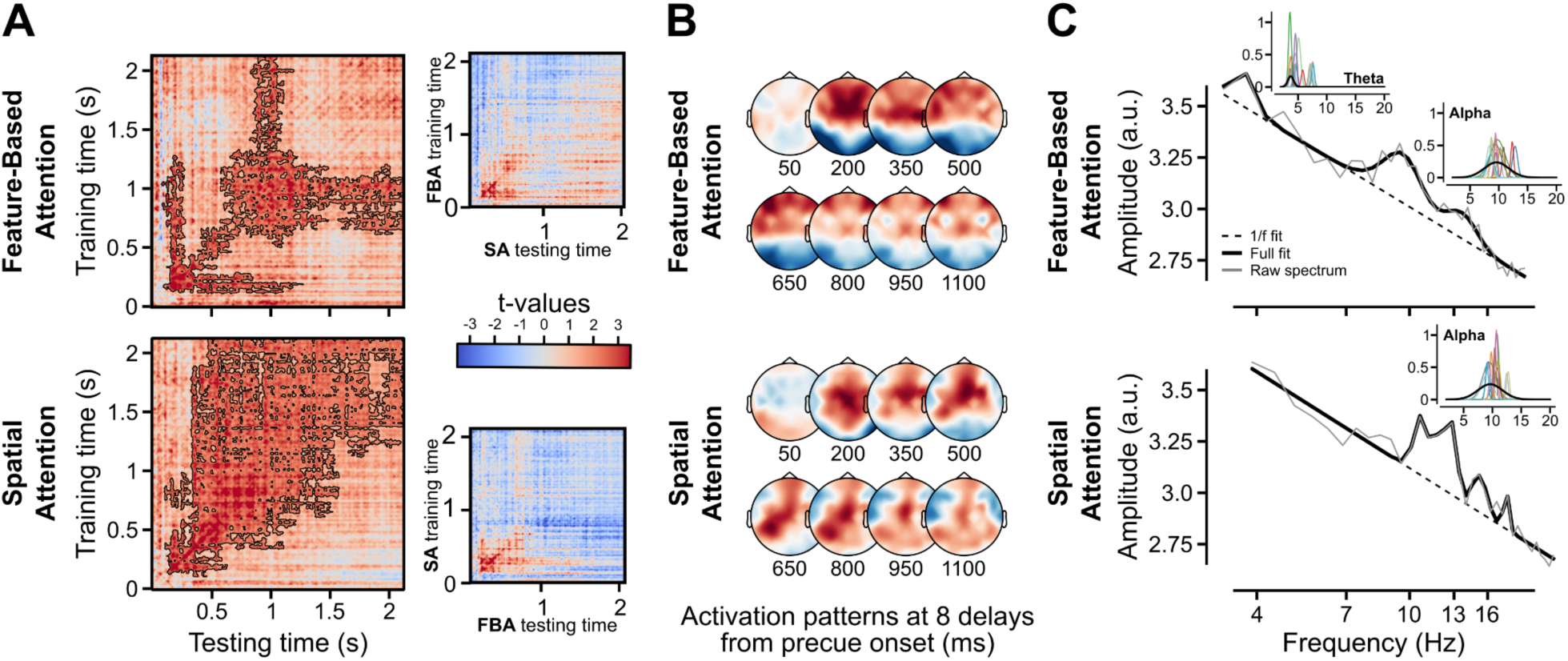
Attended feature classifications for EEG signal during attentional orienting. **A. Left** Temporal generalization for FBA (top) and SA (bottom). Diagonal represents the same training and testing time point. **Right.** Cross-condition-temporal generalization, classification on precue type (diamond vs. square). Colors represent t-values comparing classification accuracy to chance level (50%). Black contours indicate a cluster of significant above-chance classification accuracy. **B.** Activation patterns: topographies used by the classifier to discriminate the two classes. **C.** 2D-Fourier transform performed on the averaged FBA and SA 2D temporal generalization classification accuracies. Insets: same analysis performed on each participant’s spectrum.

Topographies of the activation patterns computed from classifier weights exhibited both condition-specific patterns, as well as common activation patterns to both SA and FBA, i.e., a fronto-central activation was present in both conditions (**Figure 2B**).

A cross-condition-temporal generalization analysis was further performed in which a classifier was trained at a time-point in the FBA condition and tested on every time point in the SA condition, and vice-versa (**Figure 2A right**). This control analysis showed that an increase in classification accuracy corresponding to precue evoked activity was constrained at early delays and did not lead to significant clusters (p>0.153). Together, these results confirm that the main classification analysis was not confounded by the shape of the precue itself (square/diamond), and indeed the result of the attended dimension.

### Dynamics of the stimulus space representations

Previous research showed a link between attention and rhythms in brain dynamics (for review^6,7,10^). Here, we asked whether and how rhythmic patterns of brain activity predict attention information content. We performed a 2D Fourier transform (2D FFT) on the temporal generalization matrices of each participant and condition to assess whether the observed patterns in the classification accuracies had particular spectral profiles, i.e., we asked whether the representation of stimulus dimension was modulated periodically. Akin to the 1D FFT, the 2D FFT transforms a 2D map (e.g., spatial-or temporal-domain) into a 2D frequency-domain map, enabling the identification and characterization of periodic patterns within the data. The real part of the 2D FFT was used to compute the azimuthal average and obtain a direction-independent 1D power spectrum for each map that summarizes the power spectrum independently of the direction of the periodic pattern (i.e., whether the pattern is repeated vertically, horizontally, or diagonally). Finally, frequency peaks were extracted using the specparam toolbox^34^ which parametrizes the power spectra while controlling for differences in the aperiodic structure. This was performed for each attention condition separately on the average spectra across participants, as well as on each participant’s spectrum (**Figure 2C**). The results show that alpha oscillations were present in the activity features relevant for classifying the attended dimension in both SA and FBA, whereas theta oscillations were present only in the FBA condition. Importantly, in the FBA condition, attention was sustained onto a precued feature (vertical or horizontal) and explored spatial locations (left or right). In the SA condition, attention was sustained onto a precued spatial location but explored the feature dimension. In other words, while alpha relates to both spatial and feature sustained attentional orienting (coherent with observing alpha in both SA and FBA conditions), theta is only observed during spatial exploration (only in the FBA condition). This result suggests that attention may explore the space at theta frequency.

### Attentional reorienting is supported by post-stimulus theta oscillations

We then focused on attentional reorienting during the post-stimulus period, and specifically, on how theta oscillations were affected by the specific attention conditions and topographical sites (frontal region-of-interest, ROI, and two occipito-parietal target and distractor ROIs; see **STAR Methods** for ROI selection).

In **Figure 3A-C**, we examined the power difference between invalid and valid trials for both SA and FBA conditions. For the frontal ROI **Figure 3A**, we observed higher theta amplitude in invalid compared to valid trials in the SA condition (cluster tests: p=0.0005; peak power difference ∼350 and ∼550 ms post-stimulus onset), but no significant difference between valid and invalid in the FBA condition, although a temporal cluster was marginally significant (p=0.0835; peak at ∼350 ms). Additionally, there was a significant difference between the SA and FBA conditions (p=0.0065; peak amplitude differences ∼350 and ∼600 ms). For both the target and distractor ROIs, we observed higher theta amplitude in invalid trials in both SA (target: p=0.0025; distractor: p=0.0020) and FBA (target: p=0.0140; distractor: p=0.0130) conditions (peak at ∼350 ms; **Figure 3B and C**). There was no significant difference between SA and FBA in either occipito-parietal ROI.

**Figure 3.**
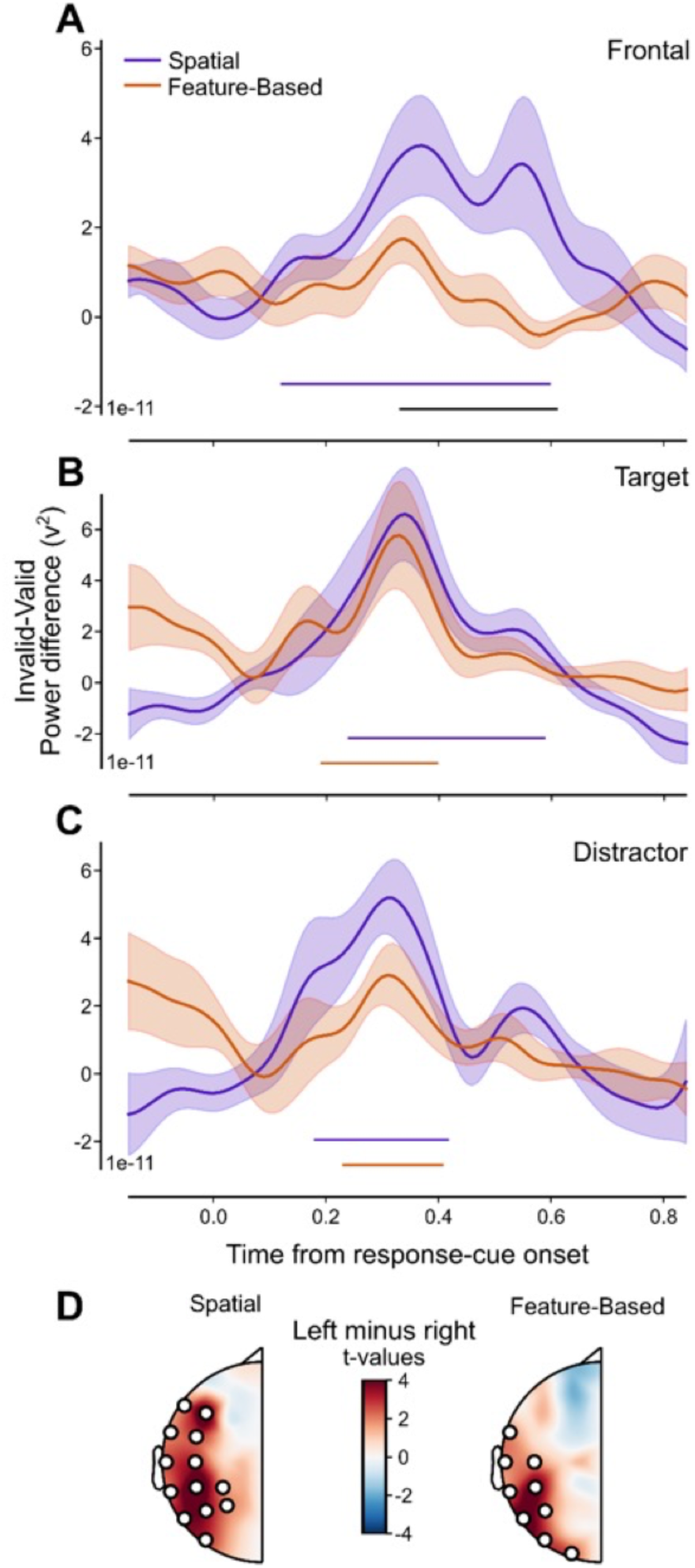
Theta oscillations during attentional reorienting. **A-C.** Theta power difference between invalid and valid trials. Theta power was estimated at each individual participant’s theta frequency peak obtained from analyses of the precue-to-stimulus period. Colored lines: significant clusters of theta power difference relative to 0 (permutation cluster test, all p-values<0.014). Black line: significant difference in theta power difference between spatial and feature-based attention conditions. **D.** Alpha lateralization maps. T-values of the target-left minus target-right difference in alpha spectral power estimates for signal extracted from 500 to 800ms from stimulus onset (positive values indicate an increase in alpha amplitude in the ipsilateral region to the target location). White dots: electrodes within a cluster of significant positive difference from zero (permutation cluster test, SA: p=0.0001; FBA: p=0.0038).

Finally, we tested whether alpha lateralization could also be observed in the late part of the post-stimulus period, i.e., after target selection^35^. Cluster-based permutation testing showed significant differences in alpha lateralization in both attention conditions (SA: p<0.0001; FBA: p=0.0038; **Figure 3D**), thus confirming that lateralization occurs when the target is spatially selected.

### Post-stimulus theta oscillations predict behavioral performance

We finally reasoned that if theta oscillations are meaningful for attentional reorienting, then they should predict behavioral performance. We used a generalized linear mixed-effects analysis with separate mixed-effects models for theta power in the 200-400 ms post-stimulus interval (interval of the highest difference between invalid and valid conditions; **Figure 3**) in each electrode ROI (frontal, target, and distractor). Each model included validity, attention conditions, and average theta power (and their interaction) as fixed effects, along with random intercepts and random slopes for validity, attention condition, and their interaction at the participant level (see **STAR Methods**). In the following paragraphs, we first present the overall effects of validity and attention condition across ROIs, then the effect of theta amplitude and its interaction with validity and attention condition per ROI. Congruent with the previous behavioral analysis (**Figure 1B**), validity was a significant predictor of RT (all p-values<0.0009) and accuracy (all p-values<0.0035) for all ROIs. The attention condition predictor only exhibited a significant effect in the model predicting RT based on theta power from the distractor ROI (χ2(1)=4.13, p=0.042). For all other analyses, the attention condition was not a significant predictor of RT nor accuracy (all p-values>0.05), which is coherent with our manipulation (staircases were used to equalize accuracy across the two attention conditions).

For all statistical results presented below, higher theta amplitude predicted better performance, i.e., higher theta amplitude predicted faster RTs and higher accuracy.

Regarding the frontal ROI, there was a significant main effect of theta amplitude on RT (χ2(1)=9.90, p=0.0017; **Figure 4A**) and accuracy (χ2(1)=5.33, p=0.021), and no interaction between frontal theta amplitude and any other factor (all p-values>0.05).

**Figure 4.**
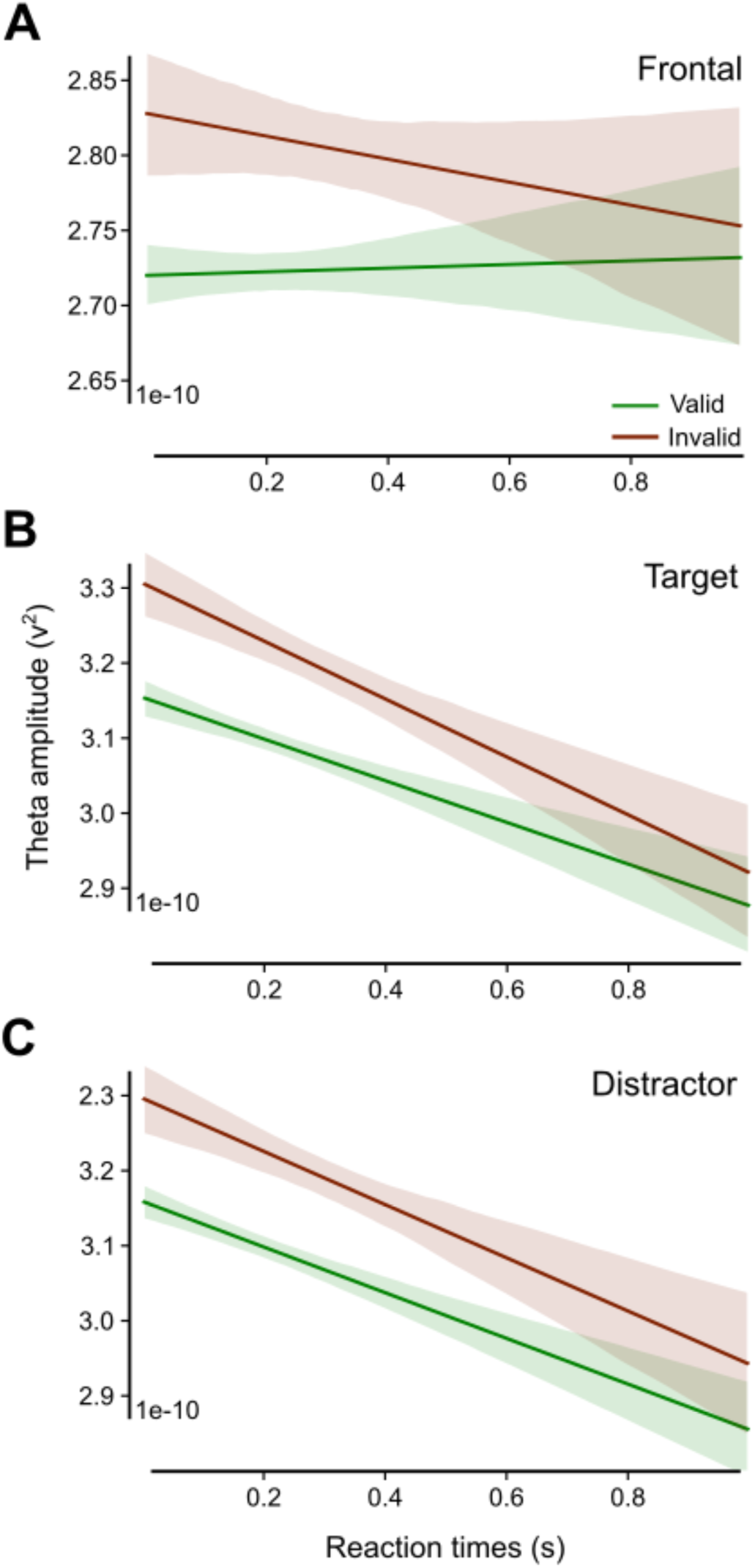
Impact of theta amplitude on reaction times. Linear regression of theta amplitude extracted at 200-400 ms post-stimulus and reaction times, for illustration purposes (independent of LME analyses). Data collapsed across attention conditions. Plots represent the regression for each validity level (green: valid; brown: invalid) for theta amplitude at the frontal (**A**), target (**B**), and distractor (**C**) ROIs.

For the target ROI, there was a significant main effect of theta amplitude on RT (χ2(1)=9.59, p=0.002; **Figure 4B**) and a trend for accuracy (χ2(1)=3.63, p=0.057). The interaction between theta amplitude and validity further showed a trend (as a predictor of RT; χ2(1)=3.77, p=0.052).

For theta oscillations in the distractor ROI, a main effect was found for theta amplitude on RT (χ2(1)=11.46, p=0.0007; **Figure 4C**) but not accuracy (χ2(1)=2.43, p=0.119), nor the interactions (all p-values>0.05). For accuracy, there was a trend for the interaction effect between theta amplitude and validity (χ2(1)=3.13, p=0.077).

Post-stimulus theta amplitude significantly predicted behavioral performance, with higher theta amplitude robustly predicting RT, and exhibiting variable predictive power on accuracy across ROIs.

## Discussion

Sustained and exploratory attention have been linked to neural oscillations in the alpha and theta frequency bands, with the sustained orienting of spatial attention associated with alpha and exploratory attention to theta (for review^7^). Our study focused specifically on clarifying the link between attentional exploration across the stimulus dimension (space/feature) and theta oscillations. Using frequency decomposition of temporal generalization matrices in the precue-to-stimulus period, we first assessed whether repeated classification patterns of brain activity predicted the specific stimulus dimension attention was oriented to. Unlike time-frequency decomposition, used to correlate oscillations with task conditions, this classification approach specifically evaluates the information content (here, the attended stimulus dimension) of repeated patterns of neural activity, directly assessing the role of such rhythmic activity in attentional orienting. The results showed alpha-rhythmic patterns of classification in both spatial (SA) and feature-based (FBA) attention conditions, suggesting that the information content was rhythmically modulated at alpha frequency with the sustained orienting of attention regardless of the stimulus dimension. Additionally, theta-rhythmic patterns were observed only in the FBA condition, i.e., when the location of the attended stimulus was unknown. This result suggests that information content of the target feature was rhythmically modulated at theta frequency when spatial exploration is made possible (see^36^ showing, consistent with our results, spatial selectivity for task-relevant locations in the prestimulus period of a feature-based attention task). We tested this hypothesis in the post-stimulus period during which we directly manipulated attentional exploration using invalid cueing. In SA, invalid cueing reoriented spatial attention from the distractor to the target location, and was previously shown to initiate theta-rhythmic attention sampling of the two locations alternatively^23,24,37^. In FBA, invalid cueing reoriented attention from the distracting stimulus feature to the other. Note, however, that in our task the two features were at two different spatial locations, which allowed us to also manipulate attentional orienting (in fact, FBA showed alpha lateralization, consistent with the now known location of the attended feature; see also^11^) and reorienting (reorienting in SA and FBA may have different temporal dynamics, but this question is beyond the scope of the current study). For both SA and FBA, the results showed that post-stimulus theta power increased in the invalid relative to the valid condition. This increase did not reach significance in the frontal ROI for the FBA. This is, however, consistent with our hypothesis. In FBA, theta oscillations were already present in the precue-to-stimulus period in both valid and invalid trials in the frontal ROI, and in the post-stimulus period, both valid and invalid trials required attentional exploration of space, albeit more in the invalid condition. Together, these would explain the reduced observed effect in this one condition. Finally, we showed that post-stimulus theta power predicted faster reaction times and higher accuracy, demonstrating its critical functional role in attentional performance. Overall, our study highlights two new findings: (1) alpha oscillations are associated with the sustained orienting of attention regardless of the attended dimension, and (2) theta oscillations support the successful spatial (and not feature) exploration by attention.

Theta oscillations have been previously related to attentional sampling and exploration of the environment using various behavioral protocols and neural measurements (for review^6,7^). In the present study, we show that theta is specifically engaged when spatial exploration is made possible (here with two possible stimulus locations). This observation challenges previous studies showing that attentional sampling of the feature dimension (with no spatial exploration) also occurs at the theta frequency^38^, and proposing a domain-general process for theta-rhythmic attentional sampling^39^. Future research will focus on combining behavioral sampling measures with EEG recordings (e.g.,^26,40^) to further assess the limits of the link between theta oscillations and attentional sampling across the stimulus dimension.

Our study further shows differential topographic patterns for attentional orienting and reorienting. In the precue-to-stimulus period, when attention orients to the cued stimulus dimension, the reported effects are observed in a fronto-central region. In the post-stimulus period, both the fronto-central and the occipital regions are involved in attentional reorienting. This observation suggests that higher-order areas are coordinating attentional allocation, and are communicating with sensory regions when stimulus processing occurs, consistent with previous proposals (for review^5–7)^. Interestingly, recent studies suggest that activity can propagate between higher-order and sensory brain regions as oscillatory traveling waves during attention tasks reflecting functional communication between regions^41,42^, and that oscillations are instrumental to content-specific top-down signals^43,44^. Future investigations will thus need to focus on characterizing the specific neural mechanisms underlying network interactions during attentional orienting and reorienting.

Together, our study clarifies the role of theta oscillations in attentional sampling. Within a single task and the same participants, we explicitly manipulated attention orienting and reorienting across the stimulus dimension. Our results suggest a restricted role of theta oscillations to spatial exploration.

### Resource availability

The custom Matlab and Python scripts are available on this repository: https://github.com/mehdisenoussi/decatsy. Anonymized data are openly available on OSF: https://osf.io/p364m/?view_only=bb19bfe5a6dd47ab8335d7536dd598ea.

## Acknowledgements

This project has received funding from the European Research Council (ERC) under the European Union’s Horizon 2020 research and innovation program (grant agreement No 852139 – Laura Dugué). We also thank Kirsten Petras, Yue Kong and Camille Fakche for their useful comments on the manuscript.

## Author contributions

M.S., N.A.B. and L.D. designed the experiment. M.S. and L.G. conducted the experiments, analyzed the data and wrote the first draft of the manuscript. All authors edited the manuscript. L.D. secured the funding. L.D. supervised the work.

## STAR Methods

### Participants

Nineteen human observers participated in all four sessions of the experiment (eight women, nine men; age [M ± SD] = 24.2 ± 5.3 years; range 21-37). Three were excluded from analysis due to technical difficulties with recording. All participants had normal or corrected-to-normal vision, gave written informed consent and were compensated for their participation. All procedures were approved by the French ethics committee Ouest IV – Nantes (IRB #20.04.16.71057) and followed the Code of Ethics of the World Medical Association (Declaration of Helsinki).

### Apparatus and Stimuli

Participants sat in a dark room, 57.5 cm from a calibrated and linearized CRT monitor (1024×786 pixels resolution, 60Hz refresh rate). A chin rest was used to stabilize head position and distance from the screen. Visual stimuli were generated and presented using MATLAB R2014 (The MathWorks, Natick, MA) and the Psychophysics toolbox version 3 (Brainard, 1997; Pelli, 1997). Display background was set to gray ([128, 128, 128] RGB). The precue and response-cue were a square (0.7°x0.7° of visual angle, [0, 0, 0] RGB) or a diamond (square rotated 90°). The stimuli were two gratings (3 cycles per degree spatial frequency; circular raised-cosine window; one horizontal and one vertical) displayed in the lower right and lower left visual quadrant. Each grating was 3° in diameter, located 3° below the horizontal meridian and 6° to the left/right from the vertical meridian. Participants were instructed to fixate a cross displayed at the center of the screen (0.6°).

### Eye-tracking

To ensure successful fixation, necessary when manipulating covert attention (with no head or eye movement), eye position was monitored throughout the experiment using an infrared video camera (EyeLink 1000 plus 5.08, SR Research, Ottawa, Canada). The start of a trial was conditioned upon fixation for 600 ms. If a fixation break occurred between precue onset and response cue offset, the trial was stopped and repeated at the end of the block (percentage of fixation breaks on average across participants (M ± SD): 14% ± 7.7, including saccades and blinks), and removed from EEG analysis. Additionally, we verified whether gaze position differed systematically between attended dimensions by comparing distance from fixation and x–y coordinates between precues, separately for each attention condition, during the precue-to–stimulus interval. No significant differences were found indicating that gaze position was comparable across precues and unlikely to have influenced the EEG results.

### Procedure

Voluntary (endogenous) Spatial Attention (SA) and Feature-Based Attention (FBA) were manipulated in two separate experimental sessions in which EEG was concurrently recorded (**Figure 1A**; session order counterbalanced across participants). Prior to each EEG session, participants performed a training psychophysics-only session on a different day.

Participants were pseudo-randomly assigned to one of four groups based on the meaning of the precues and the order of conditions. For example, participants in Group 1 underwent two sessions under the SA condition first in which the square-precue instructed them to pay attention to the bottom right quadrant, while the diamond-precue indicated the bottom left quadrant. The next two sessions were under the FBA condition, with the square-precue instructing them to pay attention to horizontal grating stimuli, and the diamond-precue indicating vertical.

*Training session.* A 4-step training session was performed the day before each EEG session. First, participants were trained to associate each precue (square or diamond) to the proper stimulus space (location or feature) using a simplified version of the trial sequence. Second, participants practiced the real trial sequence (see details in the next section) but with slower event timings and a tilt of the grating (clockwise or counterclockwise) clearly discriminable to all (10° from the horizontal or vertical). Third, a one-up two-down staircase procedure was used to adjust the tilt of the gratings to reach about 70% of detection performance (128 trials; initial tilt: 7°; step size: 3°, divided by 2 every second reversal, starting on the second reversal; minimum tilt step size: 0.1°; maximum final tilt: 30° and minimum: 0.5°; final tilt chosen from the average of the last 10 tilts). Here, the precue was replaced by a neutral (circle) cue. Fourth, participants trained on the main task using the tilt determined by the staircase.

*Main EEG experiment.* Trials started with 600 ms of successful central fixation followed by a precue displayed for 120 ms. After a 2000 ms-Inter-Stimulus Interval (precue-to-stimulus period), two grating stimuli were presented, one vertical and the other one horizontal (side randomly assigned). A simultaneously presented response-cue indicated which of the two stimuli was the target. While the grating stimuli remained on the screen for 60 ms, the response cue disappeared only after 900 ms (post-stimulus period). Participants were then asked (when the fixation cross turned black) to perform a 2-Alternative Forced-Choice (2-AFC) orientation discrimination task in which they reported the orientation of the target grating (clockwise or counter-clockwise from the horizontal or vertical meridian). If the precue and the response-cue matched, it was a valid trial (75% of trials), otherwise it was an invalid trial (25%) requiring attentional reorienting. Participants had 1000 ms to respond using the keyboard. Feedback was provided (fixation cross turning green for correct or red for incorrect responses). The next trial started after a random inter-trial interval (from 400 ms to 1400 ms). Note that the two tasks (SA and FBA) were of comparable difficulty. The average tilts of the grating in SA (mean ± standard deviation) [1.52° ± 1.38°] and FBA [1.43° ± 0.86°] conditions were not significantly different (paired t-test: t(15)=0.41, p=0.68). Each EEG session consisted of 12 blocks of 64 trials (plus repeated trials in the event of a fixation break; see **Eye-tracking** section).

### EEG Analysis

*EEG pre-processing.* EEG analysis was performed using the MNE Python toolbox version 1.6^45^ and custom scripts in Python. EEG was recorded using a Brain Products actiChamp system with 60 active scalp electrodes positioned according to the standard international 10–20 system at a sampling rate of 1000 Hz. Two electro-oculographic channels were used to record eye movements and were placed on the outer canthi of the eyes. The EEG signal was downsampled to 200 Hz and re-referenced to the average reference. Noisy electrodes (high impedance) were interpolated using a spherical spline interpolation. A notch filter at 50 Hz with a transition bandwidth of 4 Hz was applied, followed by a lowpass filter at 30 Hz. The signal was epoched from –600 ms to 3100 ms relative to precue onset, and baseline corrected using the signal between –200 ms and –10 ms. EEG and electro-oculographic epochs were visually inspected to reject visible artifacts (percentage of rejected trials across participants (M ± SD): 4.2% ± 3.0).

*Alpha lateralization.* A multitaper analysis with a Hanning window was used to estimate spectral power in the alpha frequency band (7 to 12 Hz^4^) from the EEG time series (650-1860 ms from precue onset, baseline corrected (z-score) using alpha amplitude from –350 to –80 ms). Half-topographies of alpha lateralization were computed as the alpha power difference for each electrode between precue to left and precue to right quadrant in the SA condition. Data was then collapsed onto the left electrodes, and midline electrodes set to zero. The half-topographies thus represent alpha lateralization irrespective of the cued side. The same procedure was applied for the FBA condition, by comparing precue to vertical and precue to horizontal stimulus (an additional analysis comparing trials in which the target ultimately appeared on the left versus on the right led to the same results). In the post-stimulus period, we similarly analyzed alpha lateralization using the same procedure as in ISI 1 on the interval 500-800 ms post-stimulus onset in order to limit contamination by stimuli– and motor-evoked activity. Only correct trials were selected, and valid and invalid trials were analyzed together. The half-topography represented the difference between trials in which the target was located on the right or the left in each condition separately.

*Classification analyses.* In all these analyses, the core principle was a time-resolved 2-way classification analyses on scalp topographies including all electrodes. These analyses were carried out on raw electrical potentials (i.e., broadband EEG signal after preprocessing). We trained and tested a linear discriminant analysis (LDA) classifier using a least-square solver combined with automatic shrinkage using the Ledoit-Wolf lemma. We used a stratified 10-fold cross-validation procedure. Both classifier and cross-validation were implemented using the Scikit-learn Python toolbox version 0.21.2^46,47^.

To increase signal-to-noise ratio, in all analyses we averaged four single-trials into one pseudo-trial^48,49^, and classification was performed on pseudo-trials. This procedure was performed 50 times (each with a different and random selection of single-trials) to minimize sampling bias. Decoding results were averaged across all re-samples.

For all classification results, we convolved the classification accuracy time-courses with a Gaussian kernel, with parameters mean (m)=0 and standard deviation (s)=1, to lessen classification accuracy noise. The Gaussian kernel was trimmed at 4*s, meaning that for each time point, three time points preceding, and following it, were affected by the convolution, i.e., a 35 ms-temporal window.

In the temporal generalization analysis (**Figure 2A left**), a classifier was trained on training data at time point t, and tested on test data at every time point t’, including t’=t. This procedure was repeated for each training time point t, yielding a 2-dimensional map of classification accuracies of training time versus testing time. This analysis assessed whether a discriminative pattern appearing at a given time point is informative for classification at other time points in the trial, e.g., because this neural activity pattern was sustained or re-appeared at different times^50^. Cross-temporal generalization (**Figure 2A right**) was performed similarly, but the classifier was trained and tested on different conditions, i.e., classifiers were trained to discriminate between each cue (square or diamond) in one session (e.g., FBA) and tested on discriminating cues in the other session (e.g., SA). This test assessed (1) the specificity of the classification results to the attention condition (i.e., spatial versus feature-based) and (2) to the temporal extent of the classification of the cue-evoked signal (square vs. diamond). Some studies have shown that eye-movements can confound MEG/EEG decoding analyses^51^. We thus tested whether the precues could be classified based on gaze position (time-resolved gaze x-y coordinates) using the same procedure as outlined above and found no significant above-chance classification of precues during the precue-to-stimulus interval. We finally used the method developed by Haufe et al. (2014) to extract activation patterns from classifiers’ weights in the precue-to-stimulus period reflecting the discriminative topographies used by the classifier for each training time point, for each attention condition separately (**Figure 2B**). This method accounts for the fact that multivariate classifiers’ weights reflect a mixture of noise and signal of interest. Topographies were flipped (i.e., symmetry relative to the vertical midline) depending on participants’ group to account for precue association to attended side/feature association. To lessen the noise, activation patterns’ time-courses were convolved with a gaussian kernel (µ=0 and σ=3). The kernel was trimmed at 4*σ, i.e., for each time point twelve time points preceding, and following it, were affected by the convolution (125 ms temporal window). Eight topographies were plotted for the first 1200 ms (steps of 150 ms) to keep the inset visible and because it encompasses the largest classification accuracies. Topographies show the T-value (against zero) computed at each sensor across participants per condition, thus representing the reliability of the activation patterns.

2D power spectra were computed using a 2D Fourier transform (2D FFT) to assess rhythms in classification accuracies. The azimuthal average was computed on the real part of the results to obtain a direction-independent 1D power spectrum for each map, independently for each condition and participant (code from https://github.com/keflavich/image_tools/tree/master/image_tools). The azimuthal average converts the 2D frequency domain map into a 1D profile (i.e., a frequency spectrum) that summarizes the amplitude across frequencies independently of the direction of the periodic pattern (e.g., whether the pattern is repeating vertically, horizontally, or diagonally). Finally, we used the specparam toolbox^34^ on the average spectra across participants per condition, as well as on each participants’ spectrum per condition (**Figure 2C**) to estimate peaks in a power spectrum while controlling for the structure of the 1/f noise which reflects a non-oscillatory component of the signal (also called aperiodic component).

*Spectral analyses of the post-stimulus period.* Theta oscillations’ amplitude in the post-stimulus period was assessed in data-driven selected regions of interest (ROI). The first two ROIs were selected based on the alpha-lateralization pattern (**Figure 1C**), corresponding to 8 electrodes in each occipital ROI (i.e., the 8 electrodes in the half-topographies and the symmetrical ones). The third ROI was selected using the activation patterns extracted from the temporal generalization analyzes, i.e., the electrode ROI exhibiting consistent activation across participants, for both SA and FBA. To do so, activation patterns were downsampled to 50 Hz and a spatio-temporal cluster test (see *Statistical procedure)* was performed (i.e., topographical distribution through time; 1000 permutations) for each attention mode separately. We then selected the cluster associated with the lowest p-value in each condition, and retained electrodes that were present in both clusters, corresponding to 21 fronto-central electrodes. Note that this test was used to identify ROI(s) that exhibited consistent patterns across participants and attentional conditions for further analysis, and not to assess their statistical significance.

The parieto-occipital and frontal ROIs having different numbers of electrodes, we finally selected electrodes at the center of the frontal ROI and further rejected electrodes CP1 and CP2 which appeared in both the parieto-occipital and frontal ROIs. This yielded a ROI of 7 fronto-central electrodes: Cz, C1, C2, FCz, FC1, FC2 and Fz, and two ROIs of 7 occipito-parietal electrodes each: PO4, PO8, P2, P4, P6, P8 and CP4, for the right, and PO3, PO7, P1, P3, P5, P7 and CP3 for the left. Activity in the occipito-parietal ROIs was then sorted trial-by-trial in target-or distractor-related ROIs. Trial-by-trial time-frequency decomposition was performed for the post-stimulus period interval from –300 ms to 950 ms post-stimulus onset using complex Morlet wavelet decomposition implemented in the MNE-Python suite^52,53^, for each electrode of each ROI. Frequencies ranged from 4 to 8 Hz (4 linearly spaced steps), with the number of cycles linearly increasing from 1.5 to 2 cycles. For each participant, the signal amplitude time-course in this post-stimulus period was extracted at the individual peak frequency identified in the classification accuracy maps (precue-to-stimulus period) using the specparams toolbox. Averaged amplitude time-courses across trials, for each electrode ROI, validity condition and attentional mode, were computed, for correct trials only. Finally, we computed the difference between invalid and valid trials, per participant and attention condition, and used cluster tests (see *Statistical procedure)* to evaluate whether a significant difference was present respective to 0 (i.e., no difference in theta-band activity between invalid and valid trials), as well as between the SA and FBA conditions, in each electrode ROI separately. We used one-sided tests as we hypothesized that theta amplitude would increase in invalid trials when attentional exploration was required (i.e., when attention needed to be reoriented^5–7^).

### Statistical procedure

*Cluster tests.* Inferential claims about all classification and spectral analyses were based on cluster-based permutation tests^54^ and reported following previous recommendations^33^. Cluster-based permutation analyses assessed the probability of observing a certain cluster size after thresholding the map at a predefined value by comparing the cluster size to a distribution of surrogate cluster sizes generated under the null hypothesis, which is that the data are not different from zero. Clusters were made of temporally (in the temporal generalization classification analysis for the precue-to-stimulus period and in the theta amplitude analysis for the post-stimulus period) or spatially (in the alpha lateralization analysis for the precue-to-stimulus period) adjacent samples exceeding a threshold defined based on a Student T-value distribution comparing classification accuracies to chance level (50%). This procedure yielded a p-value per cluster corresponding to the probability of observing a cluster of classification accuracies of a certain size while correcting for multiple comparisons. We used the permutation_cluster_1samp_test of the MNE toolbox v.1.6^53^, with a Hat variance adjustment, with parameter s=10^-3^, to correct for small pixel variances^55^. Surrogate cluster size distributions were generated for each participant and conditions independently by randomly permuting the sign of centered classification accuracy values, i.e., classification accuracies minus chance level, in a random subset of participants. This procedure was repeated 10,000 times for each analysis and yielded cluster p-values. From these analyses, p-values of <0.05 are depicted as black contour lines in 2D maps of classification accuracies (**Figure 2A**) and as horizontal lines in spectral analyses (**Figure 3A-C**).

*Predicting behavior from theta amplitude.* We used linear mixed effects models implemented in the lme4 package in R^50^ to evaluate whether theta oscillations in the post-stimulus period predict participants’ performance. We considered three predictors: theta amplitude, validity and attentional condition, as well as their interactions, on accuracy and RT. Models’ structure was determined using the “maximal random effects structure” approach^56^ and reduced until a significant loss in the goodness-of-fit was detected^57^.

The RT models used Gamma distributions (log link) and the accuracy models used Binomial distributions (probit link). The model structure for accuracy was

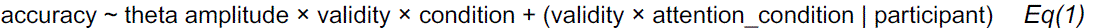

and for RT

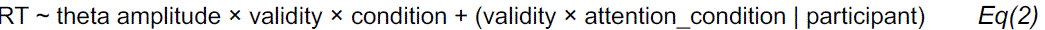

The attention_condition parameter refers to the SA or FBA condition.

